# Evidence that pain sensitivity is rhythmic in humans, mainly driven by the endogenous circadian system and little by sleep

**DOI:** 10.1101/2020.12.23.424196

**Authors:** I Daguet, V Raverot, D Bouhassira, C Gronfier

## Abstract

Pain intensity has been reported to fluctuate during the day in some experimental and clinical conditions, but the mechanisms underlying these fluctuations are unknown. Although the circadian timing system is known to regulate a wide range of physiological functions, its implication in pain regulation is unknown. We show here, using highly controlled laboratory constant routine conditions, that pain sensitivity is rhythmic over the 24-hours and strongly controlled by the endogenous circadian timing system. We find that pain sensitivity follows a sinusoidal circadian rhythmicity, with a maximum in the middle of the night and a minimum in the afternoon. We also find a weak homeostatic control of pain sensitivity, with a linear increase over the 34 hours of prolonged wakefulness, which parallels that of sleep pressure. Using mathematical modelling, we describe that the circadian system accounts for 80% of the full magnitude of pain sensitivity over the 24 hours, and that sleep-related processes account for only 20%. This result reveals that nocturnal analgesia is predominantly induced by the circadian system and has been wrongly attributed only to sleep. Our findings highlight the need to consider the time of day in pain assessment, and suggest that personalized circadian medicine may be a promising approach to pain management.

**Significance statement:** We discovered that sensitivity to pain is rhythmic in healthy humans, that sensitivity is maximal at night and minimal in the afternoon. Contrarily to the current thinking that sleep is the best painkiller, we find that the 24-h rhythmicity of sensitivity to pain is mainly controlled by a biological circadian clock in our body, and very little by our sleep. Our article reveals the neurobiological mechanisms involved in driving the rhythmicity of pain perception in humans, with the main time-piece located in the brain (the suprachiasmatic nuclei in the hypothalamus). Our findings challenge the current vision of pain physiology, and reveal the need to consider time-of-day and internal biological time for pain evaluation and pain management.

## Introduction

Pain intensity has been reported to fluctuate during the day in a number of clinical conditions (3). The cyclic nature of some headaches (4, 5) and the diurnal variation of pain related to osteoarthritis are classical clinical observations (6, 7). The mechanisms underlying these fluctuations, however, are unknown. In particular, it remains unclear whether such daily variations are related to the internal circadian clock, or to behavioral or environmental factors, such as the sleep/wake cycle or the rest-activity cycle.

Pain has two main interconnected components: a sensory-discriminative component (location, quality, duration, intensity etc.) and an emotional component (unpleasantness, anxiety, motivation, etc.) (8). This multidimensional nociceptive response involves the activation of numerous subcortical and cortical regions of the brain (e.g. somatosensory cortices, insula, thalamus, prefrontal cortex), often referred to as the “pain matrix” (9). These structures are known to be regulated by the sleep/wake cycle or the circadian clock (10–12), but it remains unclear whether pain sensitivity is rhythmic and how it is regulated.

The circadian timekeeping system plays a key role in physiology by regulating the rhythmicity of numerous functions, from gene expression to cortical activity and behavioral functions (10, 11, 13–16). It is, therefore, also likely to be involved in pain perception. The surprising lack of knowledge about the rhythmicity of pain sensitivity may result from the impact of timing on pain perception rarely having been taken into account (3), and the use of inappropriate protocols for the exploration of pain rhythmicity from a neurobiological and mechanistic point of view. The experimental studies performed to date to investigate pain sensitivity changes during the day in healthy individuals have reported conflicting results (17–22). In both experimental and clinical studies, the limited number of measurements and their timing (mostly during the daytime) made it impossible to demonstrate unequivocally the existence of a 24-hour rhythmicity in pain sensation. It is also impossible to determine the origin of any rhythmicity in pain from these studies, because neither of the two types of highly controlled laboratory protocols (constant-routine and forced-desynchrony paradigms) capable of separating endogenous and exogenous rhythms (23, 24) were used. Endogenous rhythms are controlled by the central biological clock located in the suprachiasmatic nuclei (SCN) of the hypothalamus, and exogenous rhythms depend on behavioral or environmental changes, such as the sleep/wake cycle, the dark/light cycle, or the rest/activity cycle. In real-life conditions, endogenous and exogenous influences are expressed simultaneously, making it impossible to attribute rhythmicity to one or the other. In this study, we aimed to determine whether sensitivity to heat pain displays rhythmicity over the 24-h day, and to assess the precise contribution of the circadian clock and sleep-related processes, by systematically assessing pain sensation and gold-standard markers of circadian rhythmicity in highly controlled constant-routine conditions.

## Results

### Pain rhythmicity is regulated by homeostatic and circadian processes

Twelve healthy men aged 22.7 ± 3.3 years (mean ± SEM) participated in a 56-hour experimental protocol (Figure 1) including a 34-hour highly controlled constant routine (CR) designed to unmask endogenous rhythmicity (enforced wakefulness, constant posture, low physical and cognitive activity, constant dim light, equicaloric snacks every hour)(23). We assessed the effect of time-of-day on pain sensitivity, by measuring heat pain every two hours during the 34 h of constant routine. In accordance with the current view that two main processes regulate sleep (25), and in agreement with studies showing that physiological functions (such as executive functions (26)) and cortical brain responses (measured by EEG (16) and fMRI (10)) are influenced by both sleep pressure and the circadian timing system, we then modeled the effect of time on pain with an additive mathematical model including a linear component (sleep-related homeostatic drive - process S) and a sinusoidal component (circadian drive - process C)(15).

**Figure 1.**
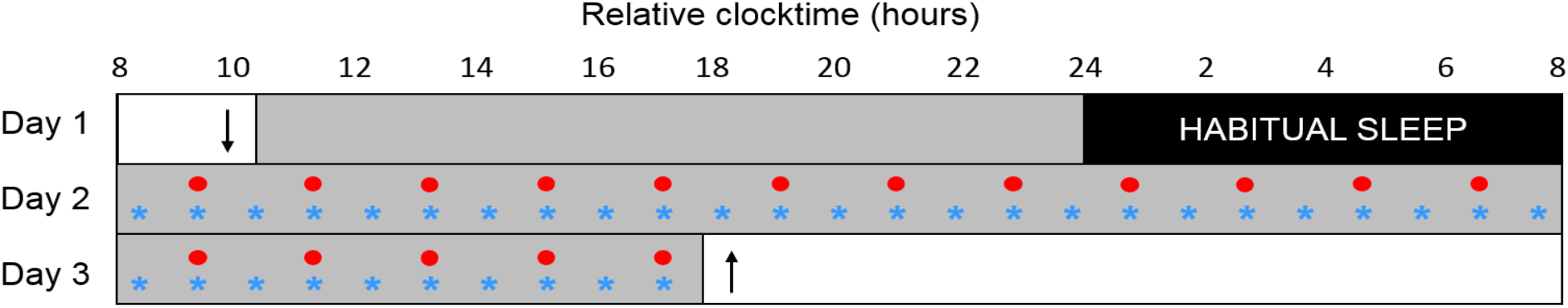
Overview of the experimental protocol. After a day of habituation (day 1) and an 8-h sleep episode, participants were subjected to a 34-hour constant routine (CR: days 2 and 3). Melatonin levels were assessed hourly (blue stars); pain sensitivity, temperature, heart rate and RMSSD were evaluated every two hours (red circles). Participants arrived at about 10:00 on day 1 (down arrow) and left the laboratory at about 18:00 on day 3 (up arrow). Gray rectangles represent wakefulness in dim light (~ 0.5 lux) and black rectangles represent scheduled sleep in darkness.

### Pain sensitivity increases with sleep debt

We probed subjective pain (visual analog scale ratings) in response to two-second heat stimuli (42 °C, 44 °C and 46 °C) every two hours over the entire 34-hour constant routine (Figure 2A, 2B and 2C; all R^2^ > 0.72). A linear component was observed for the stimuli at 44 °C and 46 °C (Figure 2E and 2F; all *p* < 0.0001; all R^2^ > 0.73), but not for the less painful stimuli at 42 °C (Figure 2D; p = 0.23; R^2^ = 0.10). Our results thus confirm the known relationship between sleep deprivation and greater pain sensitivity (27–29), but suggest that this relationship may not apply to low levels of pain. As the participants were in a constant state of wakefulness during the CR, the linear component of our model translates the effect of sleep debt and reflects homeostatic sleep pressure. The slope of the linear regression line increased with stimulation temperature (Supplementary Figure 1), so the largest changes in amplitude were observed for stimuli at 46 °C, which caused a change in pain level of 1/10 on the visual analog scale (VAS). As pain responses were measured at three arbitrary temperatures, we then used a modeling approach (classically used in pharmacology and photobiology) to extract an overall pain sensitivity value (Figure 3). The mathematically modelled sigmoidal intensity response curve (based on the combined results obtained at 42, 44 and 46 °C) yielded sensitivity values (ET_50_) that confirmed the results reported above; a linear increase in sensitivity to pain with time awake (lower ET_50_ values) (Figure 3C; R^2^ = 0.81; *p* < 0.01).

**Figure 2.**
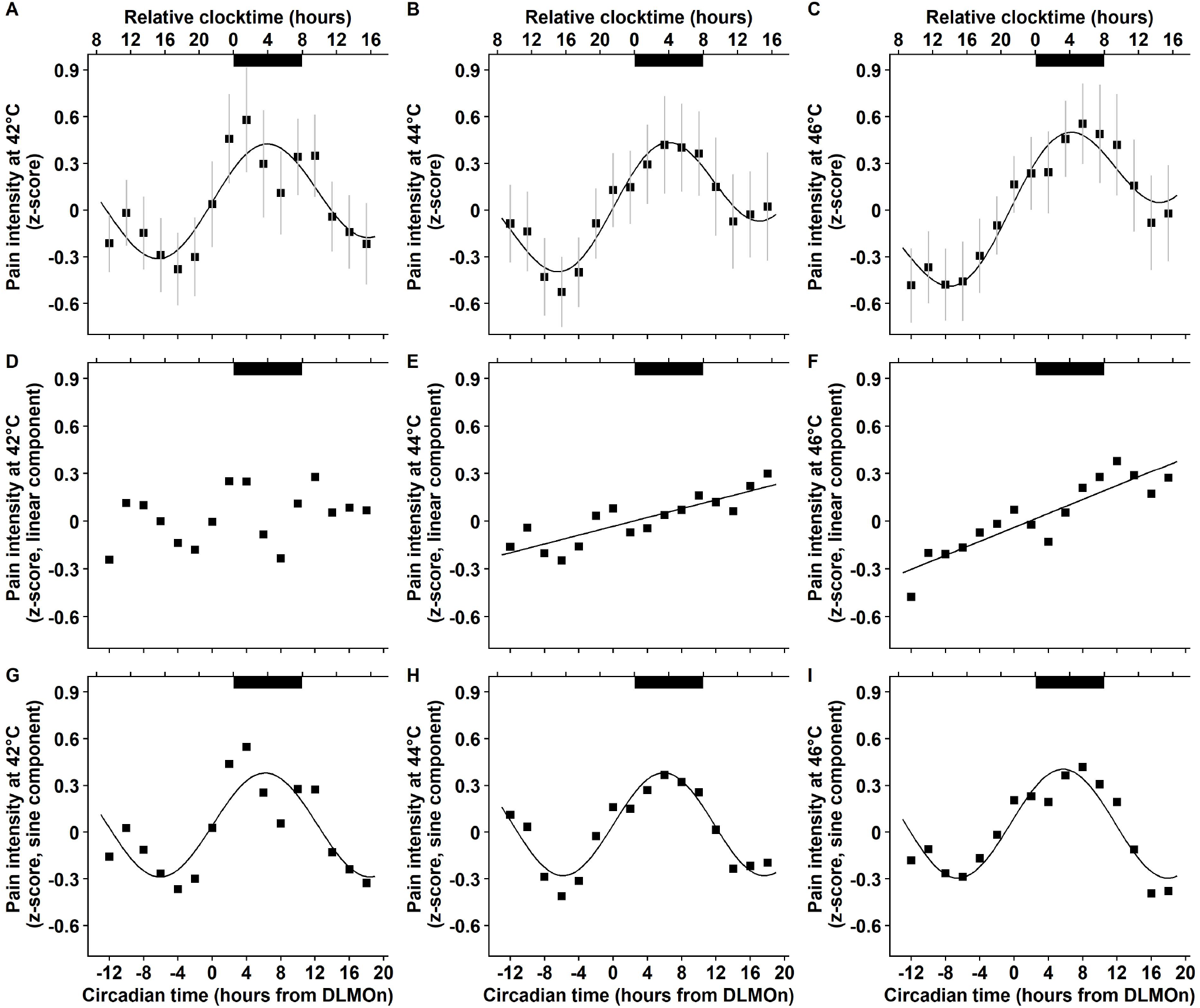
Mean pain intensities in response to 2-second heat stimuli at 42 °C, 44 °C and 46 °C are rhythmic across the 34-h constant routine protocol (*n* = 12). Dark bars correspond to the average timing of habitual sleep episodes (biological night). Circadian time 0 corresponds to dim light melatonin onset (DLMOn, mean ≃ 21:30). **A-C.** Combined models (sum of linear and sinusoidal components) applied to normalized data (mean ± SEM) for stimuli at 42 °C (**A.** R^2^ = 0.72), 44 °C (**B.** R^2^ = 0.92) and 46 °C (**C.** R^2^ = 0.92). **D-F.** Linear components for stimuli at 42 °C (**D.** R^2^ = 0.10; *p* = 0.23), 44 °C (**E.** R^2^ = 0.73; *p* < 0.0001) and 46 °C (**F.** R^2^ = 0.81; *p* < 0.00001). Pain sensitivity increases with time spent awake for stimuli at 44 °C and 46 °C. **G-I.** Sinusoidal components for stimuli at 42 °C (**G.** R^2^ = 0.70), 44 °C (**H.** R^2^ = 0.90) and 46 °C (**I.** R^2^ = 0.86). Pain sensitivity follows a circadian rhythm, with maximal pain at 3:30 (42 °C and 44 °C) or 3:00 (46 °C).

**Figure 3.**
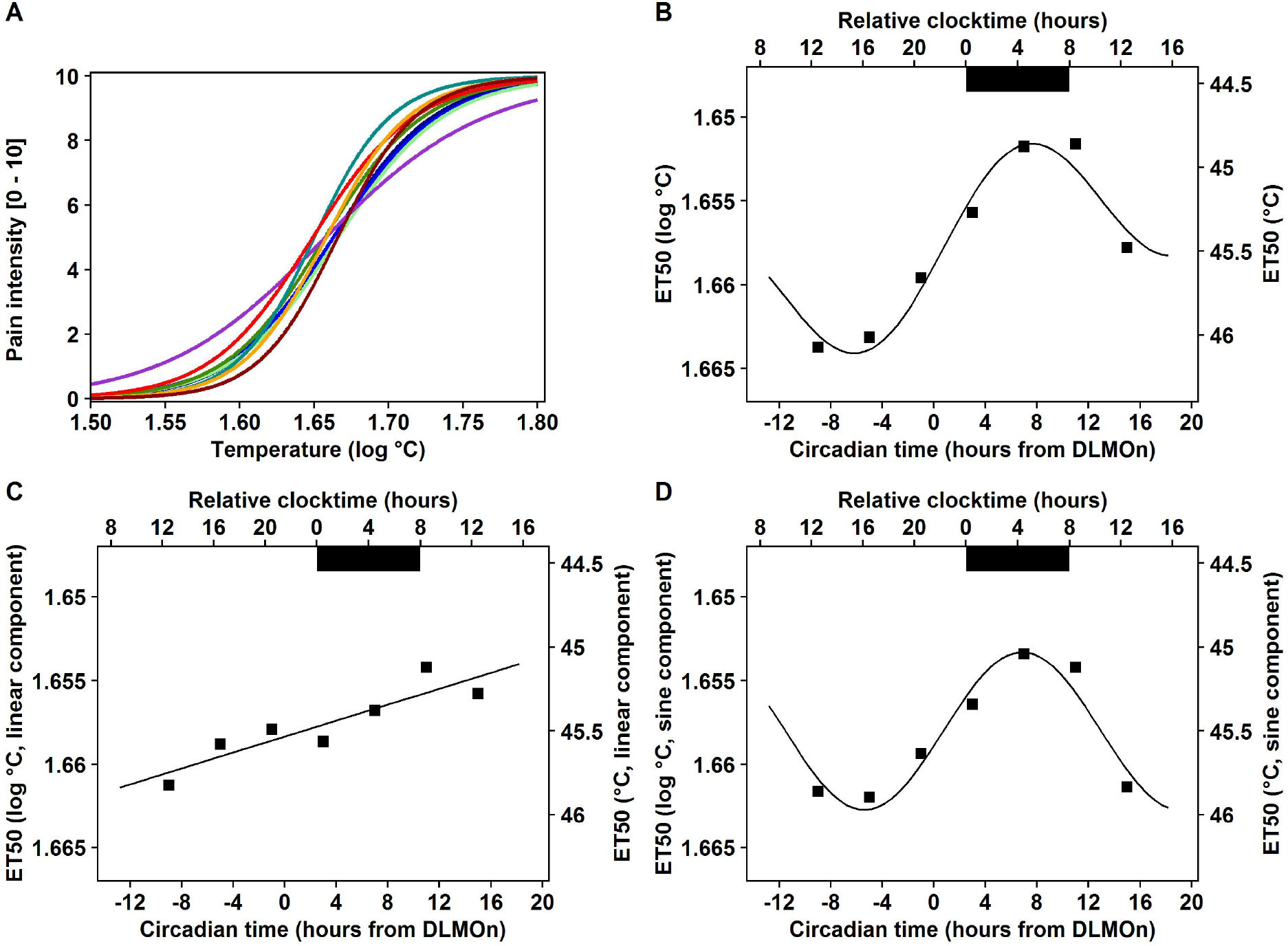
Mean pain sensitivity (ET_50_) is rhythmic across the 34-h constant routine protocol (n = 12). **A.** Intensity response curves calculated on the 6 measures obtained at 42, 44, and 46 °C over two consecutive 2-hour segments (9 curves; all R^2^ between 0.68 and 0.99). **B.** Combined model (sum of linear and sinusoidal components) applied to raw ET_50_ values (R^2^ = 0.96). **C.** Linear component (R^2^ = 0.81; p < 0.01). ET_50s_ decrease and pain sensitivity increases with time spent awake. **D.** Sinusoidal component (R^2^ = 0.93). Pain sensitivity follows a circadian rhythm with maximal pain at 4:30. **B, C and D.** Dark bars correspond to average habitual sleep episodes (biological night). Circadian time 0 corresponds to DLMOn (mean DLMOn ≃ 21:30).

### Pain sensitivity is driven by the circadian timing system, with maximal pain felt during the night

Subjective measurements of pain in response to two-second thermal stimuli (42 °C, 44 °C and 46 °C) revealed that pain sensitivity was influenced not only by sleep pressure, but also by the circadian timing system (Figure 2A, 2B and 2C; all R^2^ > 0.72). Indeed, independently of the effect of sleep pressure, a sinusoidal component in our model strongly accounted for changes in pain sensitivity across the 34 hours of constant routine, with a pain sensitivity peak between 3:00 and 4:30 for both the responses to graded stimuli (Figure 2G, 2H and 2I) and heat pain thresholds (Supplementary Figure 2C). These results were confirmed by the modeling of a sigmoidal intensity response curve, which also showed a strong circadian rhythmicity of pain sensitivity (Figure 3D; R^2^ = 0.93) and a pain peak in the middle of the night (at 4:30). Interestingly, the lack of circadian rhythmicity for warm non-painful stimuli (Supplementary Figure 3C; R^2^ = 0.13) suggests that the rhythmicity of pain sensitivity is specific to pain and is not related to a general rhythmicity of thermal sensitivity. These results provide the first evidence, to our knowledge, of a circadian rhythmicity of pain sensitivity in humans.

### Changes in pain sensitivity over the 24-hour day are mostly induced by the circadian system rather than a lack of sleep

We investigated the relative contributions of sleep and circadian drives to pain sensitivity, by calculating the mean changes in both these components and expressing them relatively to the total amplitude over 24 hours (Supplementary Figure 4). We found that the circadian system accounted for 80 % of the full magnitude of pain sensitivity changes over 24 hours, the remaining 20 % being accounted for by the homeostatic component. Surprisingly, the decrease in pain sensitivity attributable to sleep at night was very small.

**Figure 4.**
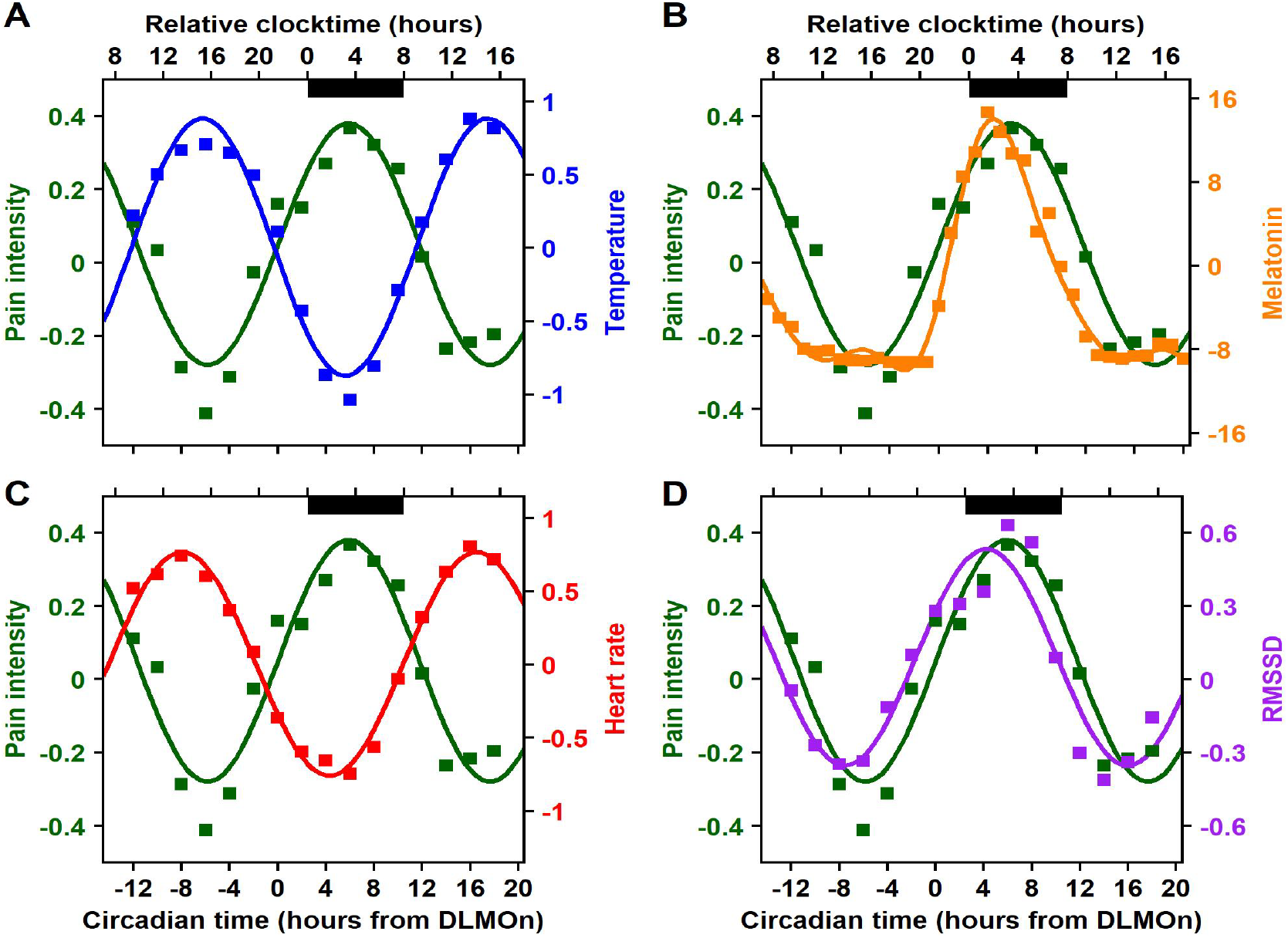
Phase relationships between circadian components of pain sensitivity and temperature (A), melatonin (B), heart rate (C) and parasympathetic activity (D) across the 34-hour constant routine protocol. Dark bars correspond to average habitual sleep episodes (biological night). Circadian time 0 corresponds to DLMOn (mean DLMOn ≃ 21:30). All curves represent the sine component of the modeled parameter. **All panels.** Circadian rhythm of VAS pain intensity scores in response to a two-second stimulation at 44 °C, with a sensitivity peak at 3:30 (green curve; R^2^ = 0.90). **A.** Circadian rhythm of baseline body temperature, with a minimal core body temperature at 3:00 (blue curve; R^2^ = 0.97). **B.** Circadian rhythm of melatonin secretion (pg/mL), with a secretion peak at 2:00 (yellow curve; R^2^ = 0 98). **C.** Circadian rhythm of heart rate, with a minimal heart rate at 2:00 (red curve; R^2^ = 0 99). **D.** Circadian rhythm of RMSDD (parasympathetic activity), with an activity peak at 2:00 (purple curve; R^2^ = 0 89).

### Phase relationships between the circadian components of pain modulation and interoceptive responses

Having identified a circadian drive for pain, we investigated whether the rhythm of pain sensitivity displayed phase relationships with interoceptive responses. Using cross-correlation analyzes, we identified a clear phase opposition (~12-h lag) between the rhythms of pain sensitivity and core body temperature (Figure 4A), with the acrophase of pain (at 3:30) occurring at about the same time as the nadir of core body temperature (at 3:00). We also found that pain sensitivity peaked 1.5 hours after endogenous melatonin secretion (at 2:00) (Figure 4B). Autonomic nervous system responses displayed strong circadian rhythmicity, with a nadir of vagal activity (minimal heart rate) and a peak of parasympathetic activity (maximal RMSSD) at 2:00, preceding the pain sensitivity peak by 1.5 hours (Figure 4C and 4D).

## Discussion

This is the first highly controlled laboratory study specifically designed to investigate pain rhythmicity and its underlying mechanisms in healthy individuals. Our results unequivocally demonstrate that pain sensitivity is endogenously driven by the circadian timing system, and that sleep and sleep deprivation have a much weaker influence on pain sensitivity than previously thought.

A limited number of peer-reviewed studies have systematically investigated the rhythmicity of pain perception in healthy individuals. Careful analysis reveals that published results are equivocal, some studies showing no rhythmicity, others reporting maximal sensitivity either during the day or during the night (17–21) A recent modelling work, using pooled datasets from four experimental studies, proposed a sinusoidal model of pain sensitivity very similar to ours, with a peak sensitivity close to midnight (43). However, because the model was built on data obtained from different populations and protocols, and collected during either sleep, wake, rest, activity, light, or dark conditions, both the phase (timing) and the origin of this rhythmicity in pain sensitivity cannot be attributed to any underlying timing mechanism, neither circadian, nor sleep-related. Overall, although often claimed by the authors, none of the previous studies we have analyzed has demonstrated that pain perception was circadian, i.e. of endogenous origin.

By contrast, our results, showing a strong sinusoidal oscillation of pain sensitivity in a constant routine protocol, i.e. in the absence of rhythmic influences and times cues, provide unequivocal evidence that the rhythmicity of pain sensitivity is driven from within, by the endogenous circadian timing system, and does not result from influences evoked the light-dark cycle, the rest-activity cycle, or the sleep-wake cycle. Indeed, if pain sensitivity were to be regulated exclusively by the sleep/wake cycle, as previously thought, we would have observed a peak in pain sensitivity at the end of our 34-hour experimental constant-routine day and not in the middle of it (after 20 hours) as we did. The very observation that the cyclicity of pain sensitivity is driven by the circadian system, independently from the sleep/wake cycle or any other environmental cycle, demonstrates that both the rhythmicity and its specific timing (its phase) are fundamental requirements in humans. Contrarily to the widely held view that pain sensitivity is driven by the sleep-wake cycle (decreasing during sleep and increasing during the day), our quantification that the circadian oscillation accounts for 80% of the full magnitude of pain sensitivity over the 24-h, and that sleep deprivation accounts for only 20% of it, reveals that sleep and sleep deprivation have in fact a very modest effect on pain. We also conclude that nocturnal analgesia is predominantly due to the circadian system, that it has wrongly been attributed only to sleep in previous studies. The pathways linking the circadian timekeeping system to pain perception cannot be inferred from this study, but the suprachiasmatic nucleus is undoubtedly the starting point, and the subcortical and cortical regions of the brain (e.g. somatosensory cortices, insula, thalamus, prefrontal cortex), often referred to as the “pain matrix” (9) are likely to be involved, given that they have been shown to be regulated by the sleep/wake cycle or the circadian clock (10–12). Our study suggests an interoceptive regulation of pain, where the circadian pacemaker is likely to be central. Pain is traditionally regarded as an exteroceptive response depending on both the somatosensory and emotional systems, however, it has been suggested that it may also be part of the interoceptive system, relating to the condition of the body (8, 44). The interoceptive responses underlying the maintenance of the internal environment of the body are organized in a hierarchical manner. They involve a number of extensively connected physiological systems, so any change in one interoceptive function is usually associated with changes in one or several other interoceptive functions. Our data are consistent with this view as they show that, like other interoceptive functions, pain is driven by a time-specific circadian rhythm that is directly related to the rhythmicity of other functions. The phase opposition we find between pain sensitivity and core body temperature (CBT) suggests an interaction between thermoregulation and nociception (45), both of which are components of the interoceptive system (8, 44). The phase relationships observed between the rhythms of the sympathetic and parasympathetic systems (as assessed by cardiovascular measurements) and pain are also consistent with this hypothesis and suggest the existence of strong interactions between the nociceptive pathways and the autonomic nervous system (ANS) (46–48). The circadian timing system may, via the SCN, serve as a key interface between pain and other interoceptive functions. The mechanisms underlying these interactions are unclear, but, interestingly, our data suggest that they are probably not mediated by melatonin. Melatonin, a nocturnal hormone released by the pineal gland, is generally reported to induce antinociceptive effects (49–51). Such effects are not consistent with the temporal relationship between peak pain sensitivity and peak endogenous melatonin secretion reported here, which instead suggests a pronociceptive effect of melatonin. None of these mechanisms can be validated on the basis of our results as we describe only temporal relationships between time series, but they could all be relatively easily tested experimentally to determine their causality. Alternatively, the circadian rhythmicity of pain may be accounted for by direct control of the nociceptive network (or the cognitive/emotional structures) by the SCN. In this regulatory model of pain regulation, the circadian system may be responsible for controlling the precise timing of nociception (8). As the thalamus is a key player in the nociceptive pathway and projections from the SCN to the anterior paraventricular thalamus have been identified (52), pain sensitivity may be directly modulated by this brain structure over the course of the 24-hour day. Multiple other pathways could be involved. Using the same highly controlled experimental conditions we employed here, a study showed that ~15% of all identified metabolites in plasma and saliva are under circadian control in humans (Dallmann 2012). These include metabolites involved in pain pathways, and recently identified metabolites of neuroinflammation specifically found elevated in patients with neuropathic pain compared to those without neuropathic pain (Pfyffer 2020). Whether those metabolites are involved in all clinical conditions of pain or in experimentally induced pain is unknown, but overlapping the human circadian metabolome and our results allows to propose that the circadian system regulates pain sensitivity though multiple pathways, both in normal and pathological situations.

The influence of sleep and sleep deprivation on pain sensitivity is modest in terms of its impact on the full magnitude of pain sensitivity over the 24-h, but it is not negligible. The linear increase in pain sensitivity that we find during enforced wakefulness, after mathematically removing the circadian component, confirms that pain sensitivity does increase with time spent awake and reveals that it is under the influence of an independent (from the circadian system) homeostatic drive, possibly related to that involved in the buildup of sleep pressure from waketime to bedtime (25). This finding is consistent with the studies we previously discussed (17–21, 43) and with the classically described interaction between pain and sleep (3, 30–33), whereby pain sensitivity appears to be driven by the sleep/wake cycle, with pain perception low in the morning after a night of good-quality sleep, increasing during the day to reach a peak before bedtime, and then decreasing during sleep (29). In the absence of sleep (after one night of total sleep deprivation), pain sensitivity has been shown to be higher than it was at the same time on the previous day (27, 28), highlighting that there is an analgesic effect of sleep and/or a hyperalgesic effect of sleep deprivation. This sleep drive is usually considered to explain why sleep disorders, such as insomnia, are associated with an exacerbation of clinical pain (29, 34). The reciprocal interactions between sleep homeostasis and pain may result from functional changes in the interconnected sleep and pain systems. Consistent with this hypothesis, sleep loss is associated with an increase in the activation of somatosensory brain areas induced by painful stimuli, potentially reflecting an amplification of neuronal responses in the cortical nociceptive systems and/or a disinhibition of normal thalamocortical pain signaling (33). In addition, sleep deprivation blunts activity in areas of the brain involved in endogenous pain modulation, such as the striatum and insular cortex (33). The specific mechanisms underlying the interactions between pain and sleep remain unknown, but may involve sleep-promoting factors, such as adenosine (35). Adenosine accumulates with increasing homeostatic sleep pressure during wakefulness, reaching high levels at the end of the day (36, 37), and then declining during sleep (38). In addition to its role in the sleep/wake cycle, adenosine is also involved in the nociceptive system and may play an anti- or pronociceptive role, depending on the receptors activated (39, 40). Thus, the hyperalgesic effect of constant wakefulness reported here may be at least partly due to adenosine accumulation, leading to A1B receptor activation (37). Obviously, other mediators, such as cytokines, which also play a role in both pain (41) and sleep regulation (42), may be involved in the sleep-related modulation of pain sensitivity.

This study has a number of limitations. First, our protocol was conducted under non-ecological and highly controlled laboratory conditions, which were nevertheless absolutely essential to dissect out the rhythmic and endogenous elements of pain sensitivity. Pain sensation may be different in real-life conditions, but the endogenous mechanisms controlling pain sensitivity are expected to be the same. The modest influence of sleep deprivation on pain sensitivity that our model finds, may also be different in real-life conditions. Indeed, prior to their experimental session in the laboratory, our participants underwent 3 weeks of sleep monitoring, during which time they slept on average 8 hours per night, and ensured they were sleep satiated upon arrival. In real life conditions, where sleep deprivation is common in our societies, the strength of sleep-related drive may be higher than in our conditions. This does not invalidate our model, but asks for its careful interpretation in different conditions (Prayag et al. 2021) and also for its evaluation in conditions of sleep deprivation. Second, pain intensity was evaluated in healthy participants, with an experimental heat pain paradigm. It is conceivable that sleep pressure and the circadian timing system have the same effect on any type of pain, but our results cannot be directly extrapolated to clinical populations. Third, the population examined in this study consisted exclusively of men. Circadian physiology is very similar in men and women, with only minor differences, such as a slightly larger amplitude (53, 54), and a slightly shorter period (55) in women, but our results should not be extrapolated to premenopausal women, in whom the menstrual cycle may modulate both the homeostatic and circadian drives of pain sensitivity.

In conclusion, our results reveal the neurobiological mechanisms driving the rhythmicity of pain perception in humans, with the main driving brain structure located in the suprachiasmatic nuclei of the hypothalamus. We show that pain sensitivity is controlled by two superimposed processes: a strong circadian component and a modest homeostatic sleep-related component. This finding may have clinical implications, as dysregulations of the circadian system have been implicated in a number of diseases with major consequences for health (13). Such alterations may also be involved in the pathophysiology of some chronic pain syndromes, as suggested for cluster headaches, for example (56). The existence of a circadian rhythmicity in pain suggests that the efficacy of pain management could be optimized using circadian medicine (1, 2). With this approach, analgesic treatments could be administered according to the each patient’s internal time (circadian time) rather than according to a uniform timing schedule mostly based on pragmatic considerations (57, 58). Such circadian approaches have already proved effective in cancer treatment (59), but have not been systematically evaluated for the treatment of pain. Individually timed medication could improve chronic pain management and greatly improve patients’ quality of life, not only by improving treatment efficacy but also by reducing the adverse effects of painkillers, including those pejorative to sleep and circadian physiology.

## Methods

### Participants

Twelve healthy men (20 - 29 years old, mean age = 22.7 ± 3.3 years; BMI = 21.8 ± 3.1 kg/m²) were included in this study. Neurological, psychiatric and sleep disorders were excluded by clinical examination and psychological questionnaires (Pittsburg Sleep Quality Index Questionnaire and Beck Depression Inventory)(60, 61). Participants had an intermediate chronotype (Horne and Ostberg Chronotype Questionnaire score between 31-69)(62) and had not done any shift work, or experienced transmeridian travel during the previous three months. Participants had normal visual acuity (Landolt Ring Test and Monoyer scale), contrast vision (Functional Acuity Contrast Test) and color vision (Farnworth D-15 and Ishihara Color Test). All experimental procedures were carried out in accordance with the Declaration of Helsinki. The study was approved by the local research ethics committee (CPP Lyon Sud-Est II) and participants provided written informed consent for participation.

### Study design

Participants were asked to maintain a regular sleep/wake schedule (bedtimes and waketimes within ± 30 minutes of self-targeted times) for an average of three weeks before admission to the laboratory, with verification by wrist activity and light exposure recordings (ActTrust, Condor Instruments, São Paulo, Brazil). Subjects were then admitted to the laboratory for a 56-hour experimental protocol (Figure 1), in which they were kept in an environment free from external time cues (clocks, television, smartphones, internet, visitors, sunlight etc.). Subjects maintained contact with staff members specifically trained to avoid communicating time-of-day information or the nature of the experimental conditions to the subjects. Participants arrived at about 10:00 on the first day. They were allowed to familiarize themselves with the laboratory environment, low light levels (< 0.5 lux), equipment, and measurements. Lunch and dinner were served at about 12:30 and 19:00. A series of measurements were then performed until bedtime (participant’s habitual bedtime), and an 8-hour sleep episode was scheduled (constant darkness; recumbent position). This was followed by a 34-hour constant-routine protocol beginning at the participant’s usual waketime on day 2, and ending on day 3 (18:00 on average). Habitual bedtimes were determined on the basis of sleep times averaged over the seven days preceding the laboratory segment of the protocol. Average bedtime was 23:45 and average waketime was 8:00.

### Constant routine protocol

A constant routine (CR) paradigm was used to reveal the endogenous circadian rhythmicity of various parameters. The CR was conducted under constant environmental conditions, to eliminate, or distribute across the circadian cycle, the physiological responses evoked by environmental or behavioral stimuli (i.e. sleeping, eating, changes in posture, light intensity variations)(23, 63). In practical terms, participants were asked to remain awake for 34 hours (starting at their habitual waketime), with minimal physical activity, while lying in a semi-recumbent (45 °) posture in bed. This posture was also maintained for the collection of urine samples and bowel movements. Room temperature (mean = 23 °C ± 0.6 (SD)) and ambient very dim halogen light levels were kept constant. Light intensity was homogeneous in the room (< 0.5 lux at the participant’s eye level in all directions of gaze). Participants were given small equicaloric snacks and fluids at hourly intervals, to maintain an equal nutritional caloric intake and stable hydration over the circadian cycle. Caloric requirements were calculated on the basis of basal metabolic rate determined with the Wilmore nomogram and were adjusted upward by a 7 % activity factor(64, 65). Fluid intake was calculated for each subject, to account for the sedentary nature of the CR(65). A member of the study staff remained in the room with the participant at all times during the CR, to monitor wakefulness and to ensure compliance with the study procedures.

### Heat and pain evaluation

Thermal stimuli were applied to the forearm with a Peltier-type thermode (30 × 35 mm) connected to a thermotest device (Somedic AB, Stockholm, Sweden). Heat detection and pain thresholds were determined according to the method of limits (mean of three measurements).

Thermode temperature was gradually increased from a baseline temperature of 32 °C, at a rate of 1 °C/s, and participants were asked to stop the increase in temperature when they started to feel a warm sensation (detection threshold) or a pain sensation (pain threshold). At this point, the temperature returned to baseline at a rate of 1 °C/s. A minimum interval of 20 s was respected between each threshold measurement. If participants had not pressed the button by the time the maximum temperature (50 °C) was reached, the stimulation was stopped and the maximum temperature was recorded as the threshold value.

The pain induced by graded thermal stimuli was assessed with a 100-mm visual analog scale (VAS). All participants received stimulation with three pseudorandomized heat stimuli (42 °C, 44 °C and 46 °C). For each stimulus, participants were asked to rate the intensity of the pain on a VAS, extending from “no pain” to “maximal imaginable pain”. For each stimulation, the thermode temperature gradually rose from baseline temperature (32 °C) at a rate of 1 °C/s. Once the target temperature was reached, it was maintained for 2 s and the temperature then returned to baseline. Stimuli were separated by an interval of at least 45 s. Pain sensitization was prevented by applying the thermode to adjacent regions of the forearm, never using the same site for consecutive stimuli.

For more precise assessments of pain sensitivity than could be achieved with the responses to arbitrary temperatures, intensity response curves were calculated (Figure 3). This is a better approach to the assessment of sensitivity, as it can be used to determine the half maximal effective temperature, or ET_50_, corresponding to the stimulation temperature required to induce 50 % of the maximal response (pain intensity of 5/10).

The data were modeled with a sigmoidal function:

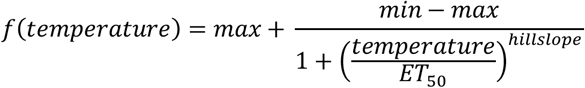

As the VAS is a bounded scale, minimum (min) and maximum (max) pain scores were set at 0 and 10, respectively. Hillslope, the slope of the curve, and ET_50_ were left free. The statistical power of the modeling approach was increased by calculating sigmoidal fits over 4-hour time epochs, corresponding to two evaluations of pain sensitivity for each of the three stimuli (42 °C, 44 °C and 46 °C), providing six points on the regression curve (Supplementary Figure 5). The ET_50_ values were extracted from each of the nine sigmoidal regressions (see formula above; Figure 3A) and plotted over time (Figure 3B).

### Body temperature

Core body temperature was measured every 2 h, with an ear thermometer (Braun Thermoscan Pro 6000, Welch Allyn, New York, USA). Body temperature was measured within 2-3 seconds, with a precision of 0.2 °C.

### Electrocardiogram

An electrocardiogram (ECG) was recorded with two adhesive skin electrodes (BlueSensor N, Ambu, Ballerup, Denmark) positioned on the sternum and the lateral thorax (RA, LL, respectively, Fontaine bipolar precordial leads). The signal was recorded at 256 Hz, with a Vitaport 4 digital recorder (Temec Instruments, Kerkrade, The Netherlands), to assess autonomic nervous system activity. Heart rate (HR) and heart-rate variability (HRV) were analyzed on the basis of the bipolar ECG signal. R-wave peak detection was performed over 10-second windows during a 4.5-minute baseline resting episode. For interval analysis, data were resampled at a rate of 10 Hz. RMSSD was determined to estimate the vagally mediated changes reflected in HRV(66, 67). It was not possible to obtain ECG data for the first participant, for technical reasons, so ECG analysis was performed for 11 participants.

### Melatonin

Saliva was collected hourly, with cotton swabs placed directly in the mouth of the participant (Salivettes, Sarstedt, Nümbrecht, Germany). Samples were stored at −20 °C until centrifugation and assay. Melatonin levels were measured with an in-house radioimmunoassay ^125^I (RIA). This assay was based on a competition technique. The radioactive signal, reflecting the amount of ^125^I-labeled melatonin, was therefore inversely proportional to the concentration of melatonin in the sample. The sensitivity of the assay was 1.5 pg/mL. The inter-assay coefficients of variation for high (18.5 pg/mL) and low (10 pg/mL) melatonin-concentration controls were 19 % and 22 % respectively, and the mean intra-assay coefficient of variation was below 10 %. We determined the circadian melatonin profile of each participant over a 24-hour day, by applying a three-harmonic regression individually to the raw data collected during the CR (days 2 and 3)(68, 69). The model equation was:

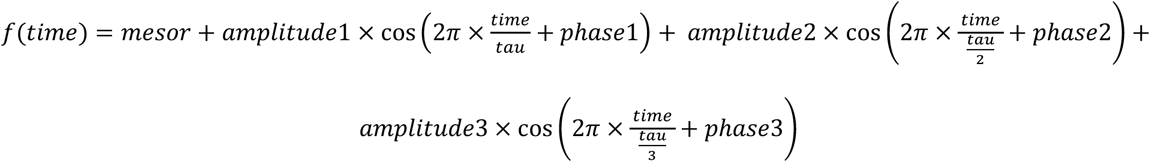

In the model, Tau (the circadian period) was constrained between 23.5 and 24.5 h; mesor, amplitudes (1 to 3) and phases (1 to 3) were set free.

The dim light melatonin onset (DLMOn), corresponding to the circadian phase, was calculated for each participant. DLMOn was defined as the time at which the ascending phase of the melatonin profile crossed the 25 % threshold of the peak-to-trough amplitude of the fitted curve. Due to technical problems with some saliva samples, the full 24-hour melatonin profile could not be obtained for two participants. For one of these participants, DLMOn was calculated on the basis of melatonin levels during the habituation day (day 1), rather than during the CR, for which we could not determine melatonin concentrations. For the second participant, in the absence of melatonin concentration data (flat profile below the limit of quantification of the assay), DLMOn was estimated from the mean phase angle calculated between habitual bedtime and DLMOn (calculated from data published by Gronfier et al., 2004)(68).

### Statistics

Outliers were identified on the basis of normalized data (*z*-scores) and were excluded from subsequent analyses (outlier.test, R, Version 3.6.1 - 2019-07-05, R Foundation for Statistical Computing, Vienna, Austria). We reduced inter-individual variability, by normalizing all data (except melatonin concentrations) by calculating individual *z*-scores and smoothing them with a moving average (calculated on 3 points). The endogenous circadian phase was taken into account for each participant, by aligning the data with the onset of melatonin secretion (DLMOn). As DLMOn occurred at different times in different participants, individual melatonin onset values were set to 0 (DLMOn = circadian time 0), and all measurement times are expressed relative to melatonin onset. We modeled the effects of time on the responses observed during the 34-hour constant routine, using an additive model including a linear component (homeostatic, process S) and a sinusoidal component (circadian, process C). The equation of the combined model was:

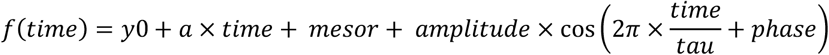

Tau (circadian period) was constrained between 23.5 and 24.5 hours(70, 71), whereas all other parameters were left free. Once the parameters of the combined model had been defined, process S and process C were modeled separately. The homeostatic component (process S) was regressed against the linear component of the model: *f*(*time*) = *y*0 + *a* × *time*. The circadian rhythmicity (process C) of the data was regressed against the sinusoidal component of the model: 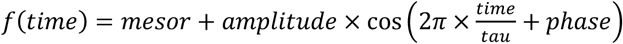.

Statistics were calculated with R (Version 3.6.1 - 2019-07-05, R Foundation for Statistical Computing, Vienna, Austria). Results were considered significant if *p* < 0.05. Unless otherwise stated, results are expressed as means ± SEM.

## Author contributions

The experiment was conceived by CG, and designed by CG, DB and ID. Data were collected by ID and CG. Melatonin assays were conducted by VR. Data were analyzed and interpreted by ID and CG. ID, CG and DB wrote the manuscript, and VR revised the manuscript. All authors agree to be accountable for all aspects of the work.

## Acknowledgments

We wish to thank all the volunteers who participated in this study. We also wish to thank the staff and students, and especially Pauline Kirchhoff who participated in data collection and analysis. Special thanks also go to Dr Alain Nicolas, who conducted the medical and physical examinations.

## Supplementary information

**Supplementary Figure 1.**
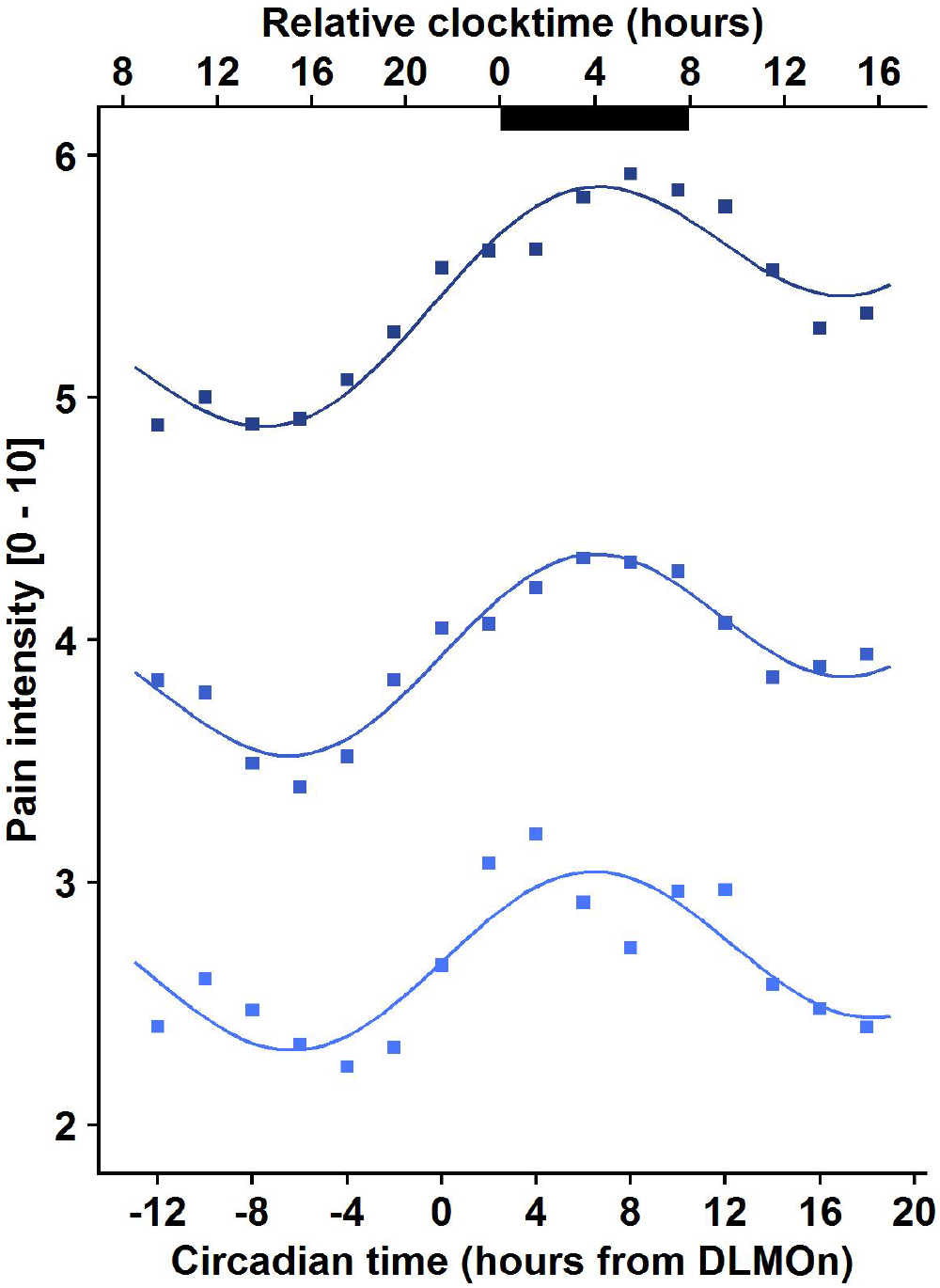
Mean pain intensity in response to heat stimuli at 42 °C, 44 °C and 46 °C, across the 34-hour constant routine protocol (n = 12). Dark bars correspond to average habitual sleep episodes (biological night). Circadian time 0 corresponds to DLMOn (mean DLMOn ≃ 21:30). Combined models (sum of linear and sinusoidal components) of pain sensitivity in response to heat stimuli at 42 °C (light blue curve), 44 °C (blue curve) and 46 °C (dark blue curve). The amplitude changes (between peak and through) in pain intensity for 42 °C, 44 °C and 46 °C are of 0.7, 0.8, and 1 VAS respectively.

**Supplementary Figure 2.**
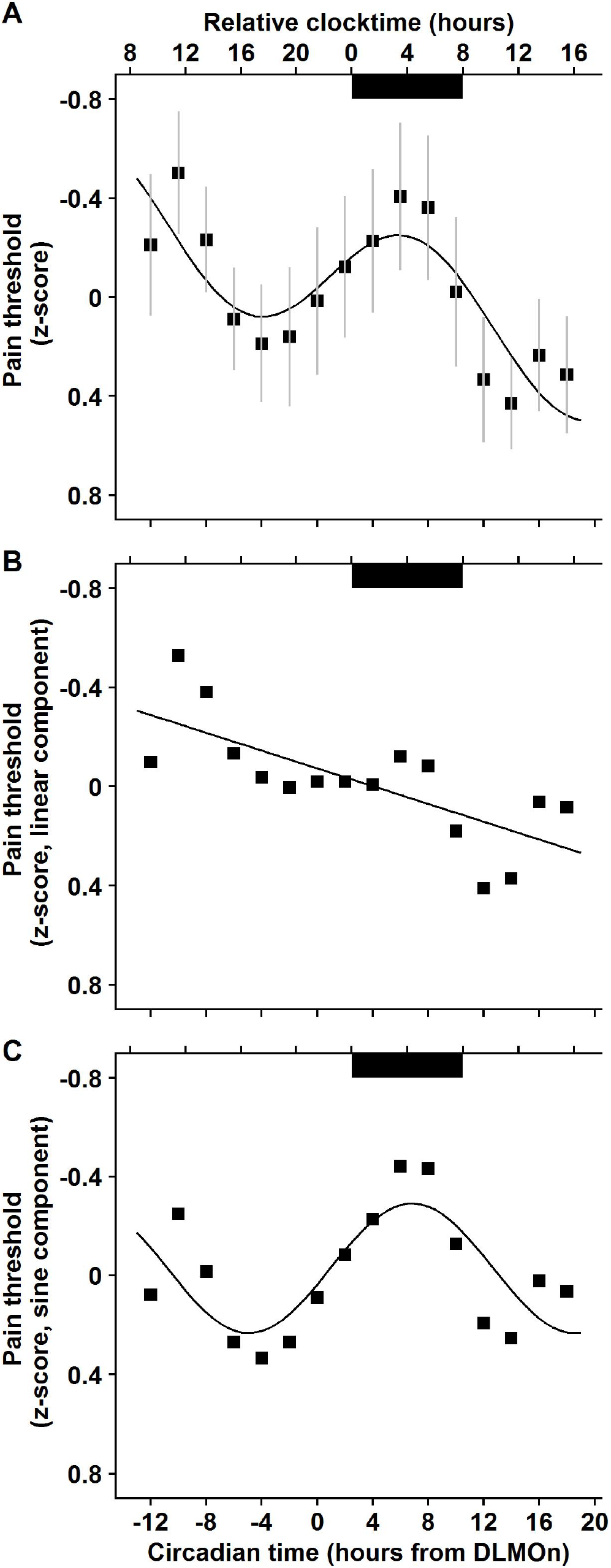
Mean heat pain thresholds, across the 34-h constant routine protocol (n = 12). Dark bars correspond to average habitual sleep episodes (biological night). Circadian time 0 corresponds to DLMOn (mean DLMOn ≃ 21:30). **A.** Combined model (sum of linear and sinusoidal components) applied to normalized data (mean ± SEM; R^2^ = 0.69). **B.** Linear component (R^2^ = 0.53; p < 0.01). Pain sensitivity decreases with time spent awake. **C.** Sinusoidal component (R^2^ = 0.57). Pain sensitivity follows a circadian rhythm with maximal pain at 4:30.

**Supplementary Figure 3.**
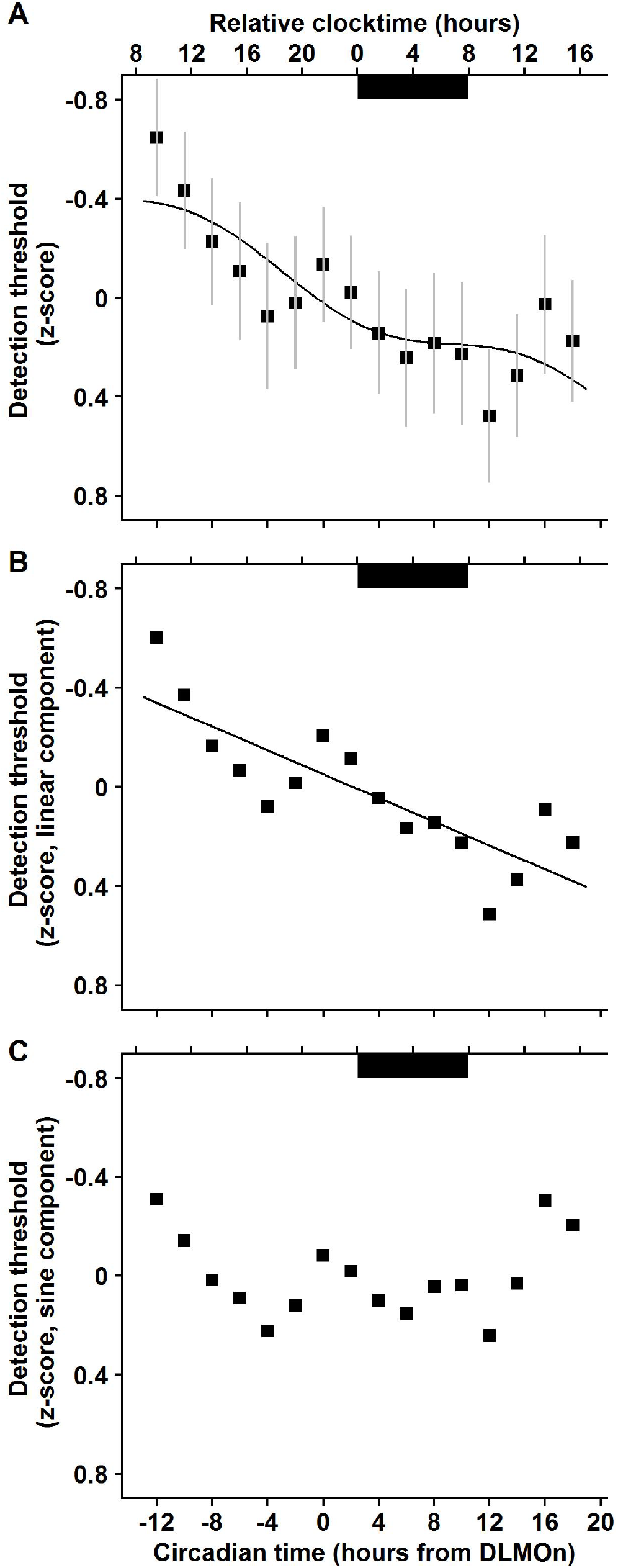
Mean warm detection thresholds across the 34-h constant routine protocol (n = 12). Dark bars correspond to average habitual sleep episodes (biological night). Circadian time 0 corresponds to DLMOn (mean DLMOn ≃ 21:30). **A.** Combined model (sum of linear and sinusoidal components) applied to normalised data (mean ± SEM; R^2^ = 0.70). **B.** Linear component (R^2^ = 0 68; p < 0.0001). Heat sensitivity decreases with time spent awake. **C.** No sinusoidal component is found (R^2^ = 0.13).

**Supplementary Figure 4.**
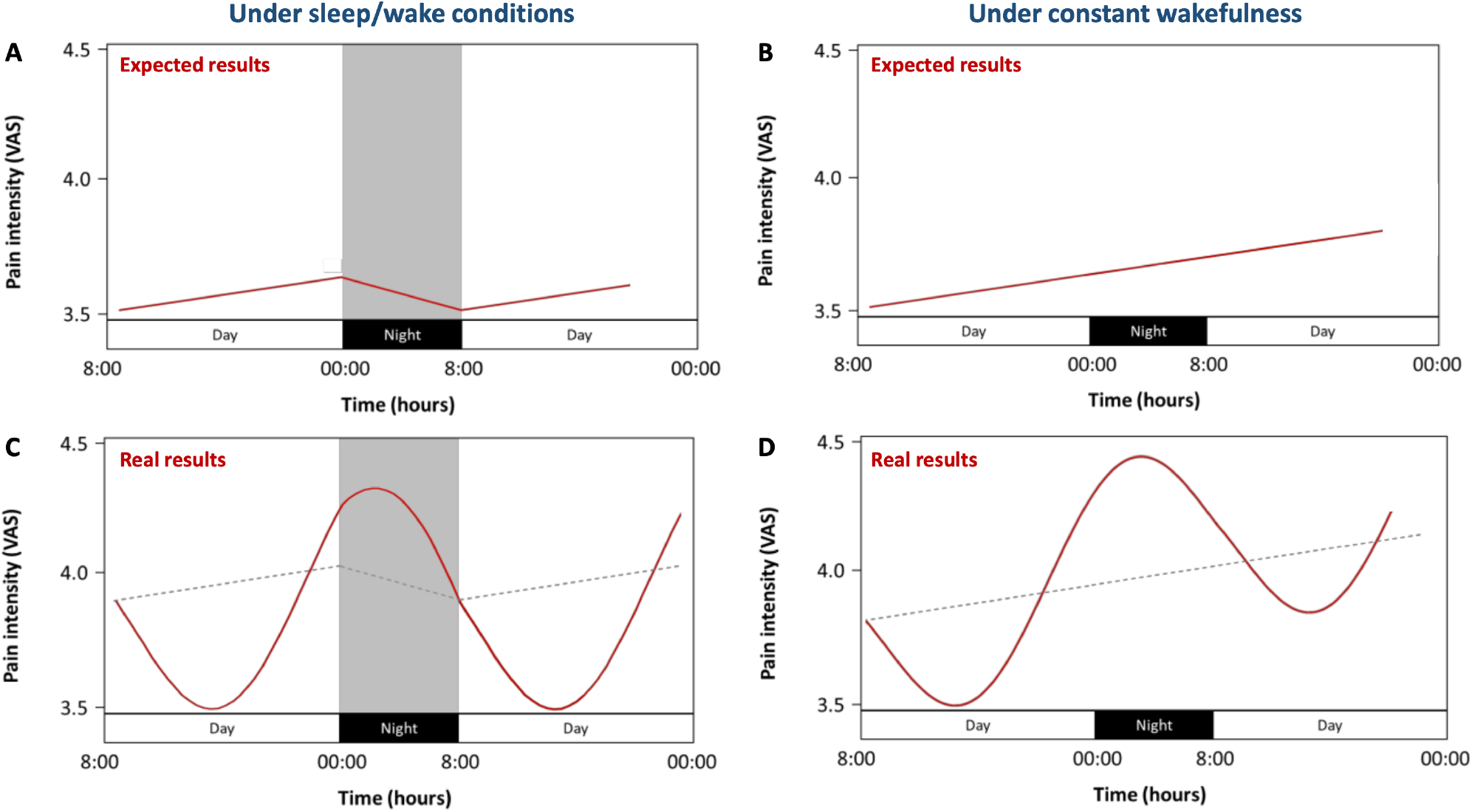
Variations of pain sensitivity across the 24h day. A-B. Expected pain sensitivity, according to the current homeostatic model based on data obtained in the literature (based on 3 articles only). Under regular sleep/wake conditions (**A**), pain sensitivity increases during wakefulness, in parallel to sleep pressure, and decreases during the night, due to an analgesic role of sleep. Under constant wakefulness conditions (**B**), pain sensitivity increases during wakefulness, and keeps increasing in the absence of sleep. In this model, pain sensitivity only depends on time since awakening (during the day), and time since bedtime (at night). **C-D.** Our results show that pain sensitivity is driven by two independent and additive components: a homeostatic drive and a circadian drive. With sleep at night (**C**) both mechanisms co-exist. Pain oscillates sinusoidally (circadian drive) and pain increases linearly during wakefulness and decreases during sleep (homeostatic drive – grey dotted line). Without sleep at night (**D),** our results under constant wakefulness show the superimposed additive homeostatic (grey dotted line) and circadian regulation of pain, with both a linear increase with time spent awake, and a sinusoidal oscillation.

**Supplementary Figure 5.**
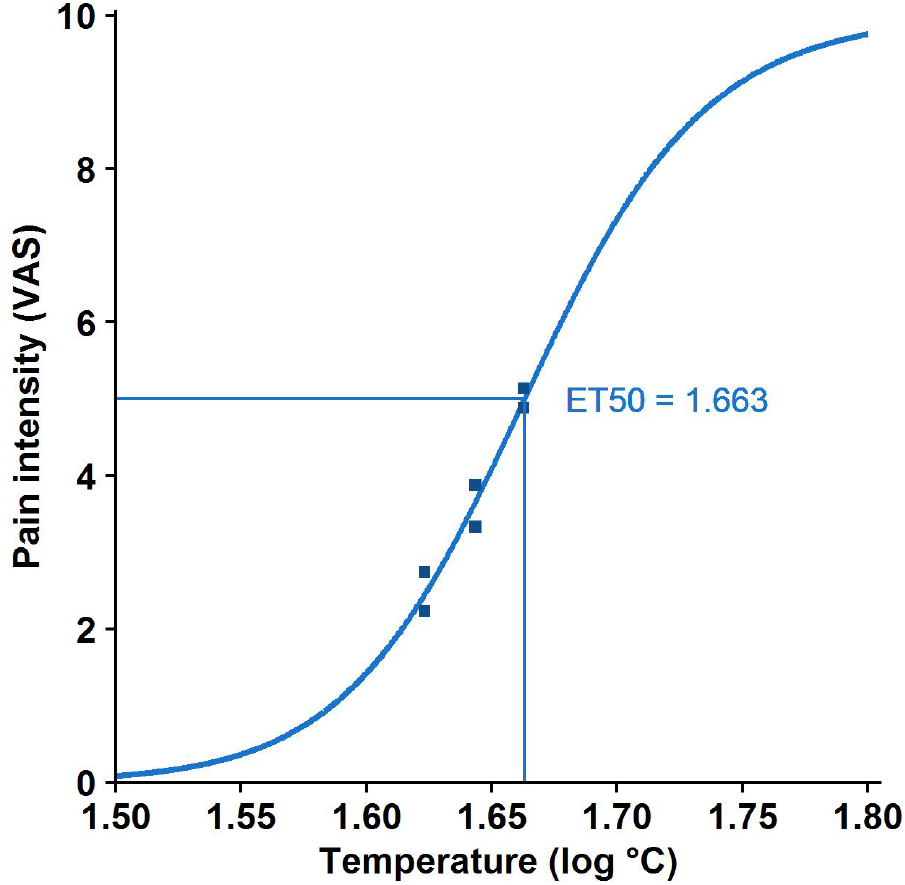
Pain Intensity Response Curve (IRC) at circadian time −9 (~ 12:30). Sigmoidal regression, calculated on a 4-hour time epoch (10:30 – 14:30), corresponding to 2 evaluations of pain sensitivity to each of the 3 heat stimulations (42 °C, 44 °C and 46 °C), providing 6 points for the regression. The ET_50_ value, corresponding to a stimulus inducing a pain of 5/10, is extracted from the sigmoidal regression (here ET_50_ = 1.66 log[temperature]).

### Time-of-day mechanisms of heat detection and pain thresholds

Heat pain thresholds and warm detection thresholds were measured every 2 hours throughout the whole 34-hour constant routine (Supplementary Figures 2 and 3; all R^2^ > 0.69). The significant linear trend observed for heat pain thresholds (Supplementary Figure 2; p < 0.01; R^2^ = 0.53), suggests a decrease in pain sensitivity with sleep pressure. This result might reflect a deterioration of cognitivo-motor performances (slower reaction times) associated with sleep pressure *(1,2)*. The fact that this effect was not specific to the heat pain threshold, since a similar relationship was seen for the warm detection threshold (Supplementary Figure 3; R^2^ = 0 68; p < 0.0001), is consistent with this hypothesis. The strong circadian rhythm of pain sensitivity (with a peak at 4:30), assessed through heat pain threshold measures, confirms the results found with graded heat stimuli (and presented in the main article). The lack of circadian rhythmicity for warm non-painful stimuli (Supplementary Figure 3C; R^2^ = 0.13) suggests that the rhythmicity of pain sensitivity is specific to pain and is not related to a general rhythmicity of thermal sensitivity.

